# Cx43 carboxyl terminal domain determines AQP4 and Cx30 endfoot organization and blood brain barrier permeability

**DOI:** 10.1101/2021.09.30.462547

**Authors:** Antonio Cibelli, Randy Stout, Aline Timmermann, Laura de Menezes, Peng Guo, Karen Maass, Gerald Seifert, Christian Steinhäuser, David C. Spray, Eliana Scemes

## Abstract

The neurovascular unit (NVU) consists of cells intrinsic to the vessel wall, the endothelial cells and pericytes, and astrocyte endfeet that surround the vessel but are separated from it by basement membrane. Endothelial cells are primarily responsible for creating and maintaining blood-brain-barrier (BBB) tightness, but astrocytes contribute to the barrier through paracrine signaling to the endothelial cells and by forming the glia limitans. Gap junctions (GJs) between astrocyte endfeet are composed of connexin 43 (Cx43) and Cx30, which form plaques between cells. GJ plaques formed of Cx43 do not diffuse laterally in the plasma membrane and thus potentially provide stable organizational features to the endfoot domain, whereas GJ plaques formed of other connexins and of Cx43 lacking a large portion of its cytoplasmic carboxyl terminus are quite mobile. In order to examine the organizational features that immobile GJs impose on the endfoot, we have used super-resolution confocal microscopy to map number and sizes of GJ plaques and aquaporin (AQP)-4 channel clusters in the perivascular endfeet of mice in which astrocyte GJs (Cx30, Cx43) were deleted or the carboxyl terminus of Cx43 was truncated. To determine if blood-brain-barrier integrity was compromised in these transgenic mice, we conducted perfusion studies under elevated hydrostatic pressure using horseradish peroxide as a molecular probe enabling detection of micro-hemorrhages in brain sections. These studies revealed that microhemorrhages were more numerous in mice lacking Cx43 or its carboxyl terminus. In perivascular domains of cerebral vessels, we found that density of Cx43 GJs was higher in the truncation mutant, while GJ size was smaller. Density of perivascular particles formed by AQP4 and its extended isoform AQP4ex was inversely related to the presence of full length Cx43, whereas the ratio of sizes of the particles of the AQP4ex isoform to total AQP4 was directly related to the presence of full length Cx43. Confocal analysis showed that Cx43 and Cx30 were substantially colocalized in astrocyte domains near vasculature of truncation mutant mice. These results showing altered distribution of some astrocyte nexus components (AQP4 and Cx30) in Cx43 null mice and in a truncation mutant, together with leakier cerebral vasculature, support the hypothesis that localization and mobility of gap junction proteins and their binding partners influences organization of astrocyte endfeet which in turn impacts BBB integrity of the NVU.

## INTRODUCTION

Astrocytes, once viewed as passive space-filling neural support cells, are now appreciated as highly polarized and specialized cells that play a major role in dynamic control of neuronal activity. Astrocyte processes extend to each of the other neural components, creating distinct functional compartments: The interfaces between individual astrocytes and heteroglial contacts with and oligodendrocytes, the cradle around the synapse, and the endfoot surrounding endothelial cells in the neurovascular unit. A remarkable feature of astrocytes is that at each of these regions, gap junctions are present and produce a pan-glial syncytium enabling long range signaling within anatomically defined areas; gap junctions in astrocytes provide heterocellular, heterotypic gap junctions with oligodendrocytes, the formation of both reflexive (or “autaptic”) junctions with itself (see ^1^) and homotypic junctions with other astrocytes ^2^. Gap junctions consist of core channel proteins, the connexins (primarily Cx43 and Cx30 in astrocytes), to which are attached scaffolding, signaling and cytoskeletal proteins, assembling into what we have termed the Nexus ^3^. Intercellular gap junction channels are the primary mechanism by which astrocytes communicate among themselves and with oligodendrocytes, and the other Nexus components likely act to modulate intercellular communication and provide the additional functions proposed for gap junctions, including adhesion, cell migration/process motility, and maintenance of both gap and tight junctions in neighboring cells.

The astrocyte endfoot serves as the gateway for delivery of water, ions and metabolites into the brain parenchymas, and location of channels and transporters for this function is highly polarized to face the perivascular space. AQP4 at the perivascular endfoot and intercellular gap junction channels connect astrocytes into a water distribution network that extends from the perivascular space throughout the parenchyma. This flux drives bulk flow of water, ions and small molecules, in a process termed glymphatic circulation that is driven by cardiovascular pulsation and is crucial to delivery of nutrients and removal of debris. Water uptake and delivery throughout brain parenchyma are highly interdependent, as evidenced by enhanced gap junction coupling resulting from AQP4 deletion ^4^. Our studies using fluorescence recovery after photobleaching (FRAP) to determine mobilities of gap junction plaques with respect to other membrane proteins have revealed that gap junctions formed of Cx43 are ordinarily highly stable, whereas paired Cx30 connexons rearrange rapidly ^5^. Truncation of the cytoplasmic terminus or mutagenesis of cytoplasmic cysteine residues converts Cx43 behavior to that of mobile Cx30 ^6^. Stability of the Cx43 plaque was also shown to dictate distribution and mobility of other membrane proteins, lipids and cytoplasmic Nexus components ^7^, leading to the hypothesis that Cx43 plaques serve as structural determinants of endfoot organization.

Consistent with this hypothesis, transgenic mice lacking astrocyte connexins display numerous abnormalities, including leaky blood-brain-barrier and edematous astrocyte endfeet with reduced AQP4 and beta-dystroglycan expression ^8^, and abnormal white matter with vacuolated oligodendrocytes ^9^. Moreover, mice have been generated in which full-length Cx43 has been replaced with Cx43 mutant truncated at amino acid 258, immediately upstream of the region we have implicated in conferring junction stability. Homozygous Cx43(K258stop) mice are not viable due to severe edema, attributed to loss of the epithelial junctional barrier ^10^; mice heterozygous for truncated Cx43 (Cx43^K258/-^) show defective migration of neuronal precursors ^11^, altered intercalated disc structure and localization ^12^ and predisposition to cardiac arrhythmias and larger infarct size after cardiac or cerebral ischemia ^13,14^. It is noteworthy that the location of this Cx43 mutation coincides with a frame-shift mutation responsible for occulodentodigital dysplasia (ODDD), a syndrome with major neurological impairments ^15,16^.

These prior studies indicate that astrocyte endfeet and the gap junctions that connect them are essential for the proper molecular organization of the neurovascular unit and for its barrier function. To test the extent to which the immobile Cx43 gap junction plaques impose structural order that enables barrier formation, we have used a model of high-pressure perfusion to evaluate whether transgenic mice with altered connexin expression or not possessing the Cx43 carboxyl terminus scaffolding-binding domain exhibit differential susceptibility of brain vasculature to rupture and have used super-resolution microscopy to examine organization of endfoot proteins. Our findings provide evidence that presence of Cx43 and its cytoplasmic binding domain are important determinants of structural aspects of AQP4 aggregates and Cx43 gap junction plaques. We propose that through its linkage to scaffolding and other Nexus components, cytoskeletal elements and endfoot markers, Cx43 provides stability and overall structural organization for the astrocyte endfoot.

## RESULTS

### 1. Distribution and expression of Cx43 and Cx30 in the different transgenic mice

In this study we have examined brains from 4 transgenic mouse lines in which astrocyte gap junction expression has been altered, as illustrated in Figure 1A. Immunostaining sections with the endothelial cell marker PECAM 1 (CD31) revealed that both Cx43 (Fig. 1B) and Cx30 (Fig. 1C) are expressed throughout the brain, with Cx43 being particularly abundant in proximity to brain vasculature. As considered in more detail below, perivascular location of these connexins overlapped to some extent.

**Figure 1.**
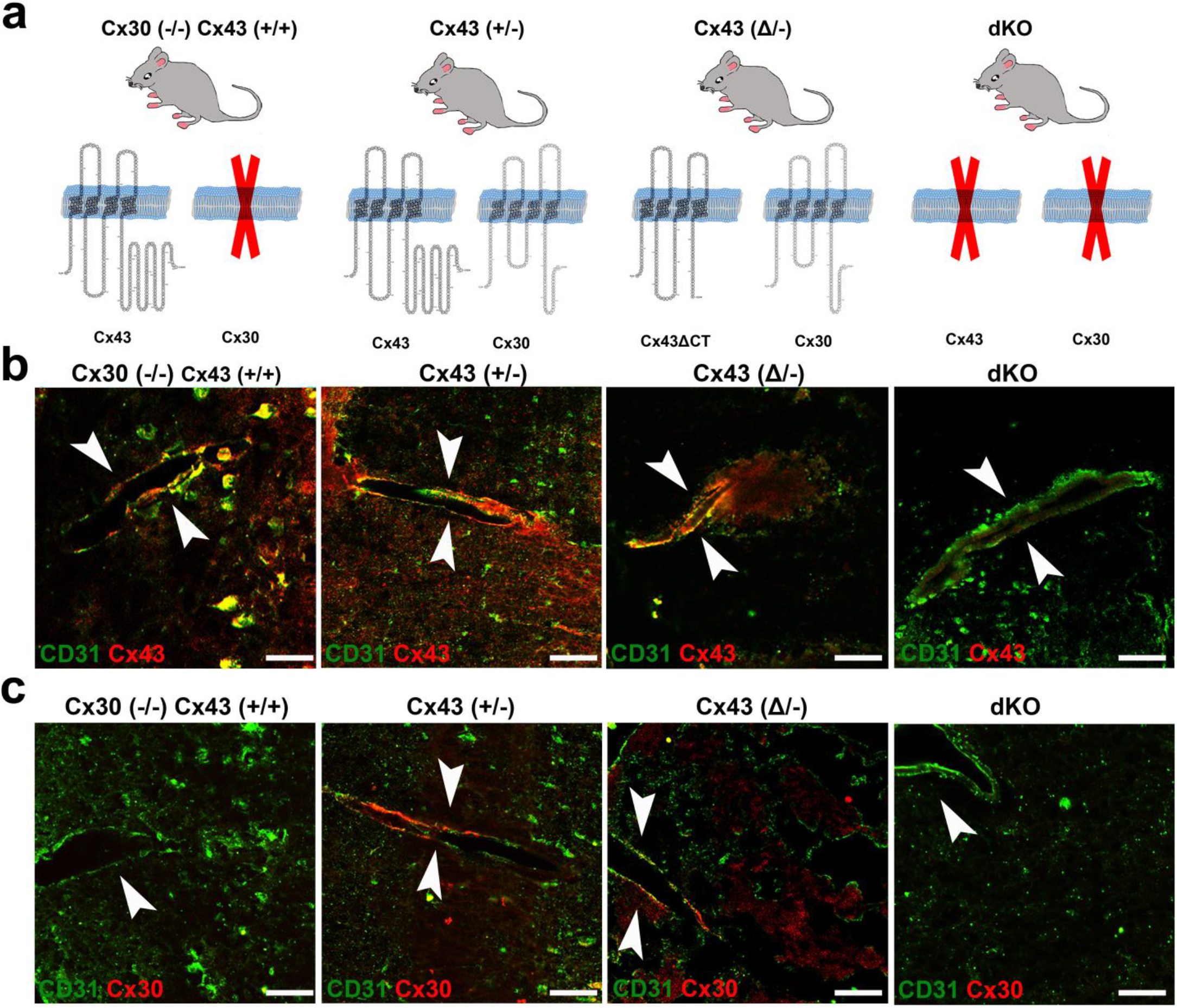
Mice used and general appearance of Cx30 and Cx43 in perivascular astrocyte domains. (A) Diagrams of gap junction composition of transgenic mice used in this study arranged in order of decreasing Cx43 expression. Mice used were: Cx43 WT but Cx30 null [Cx43^(+/+)^/Cx30^(-/-)^], Cx43 heterozygous [Cx43^(+/-)^/Cx30^(+/+)^], heterozygous for the M258 truncation [Cx43^(□/-)^/Cx30^(+/+)^], and dKO [hGFAP-Cre:Cx43^(fl/fl)^/Cx30^(-/-)^]. (B, C) Double immunolabeling with CD31 to localize endothelial cells in brain vessels and with antibody for either Cx43 (B) or Cx30 (C) in each of the genotypes illustrated in Note localization of both Cx30 and Cx43 nearby CD31, indicating that both connexins are present at astrocyte endfeet. Arrowheads indicate the vessels. Scale bar: 50µm.

### 2. Leakiness of the blood brain barrier in mice with astrocytes lacking Cx43 or its carboxyl terminal domain

Previous studies reported the observation that the volume of tissue damage after middle cerebral artery occlusion (MCAO) ischemic stroke model was greater in the Cx43^(Δ/-)^ mouse ^13^ and that in the dKO mouse brain high pressure perfusion (HPP) led to higher vascular leakage of HRP than in the wildtype mouse ^8^. However, the extent of the difference was not quantified. To explore the relation between Cx43 expression and blood brain barrier permeability in a greater detail, we quantified the impact of both Cx43 truncation and deletion using the HPP model. As illustrated in Fig. 2A, microhemorrhages were readily detectable in brains of mice of all genotypes after high pressure perfusion. Whereas density of infarcted regions did not differ between Cx43^(+/+)/^Cx30^(-/-)^ and Cx43^(+/-)^ brains, the number of micro-hemorrhages per area was higher for both Cx43^(Δ/-)^ and dKO mutants compared to their respective controls (Fig. 2B). With regard to the sizes of infarcts, areas were larger in the Cx43^(+/-)^/Cx30^(+/+)^ and lowest in dKO brains (areas ∼2000-4000 μm^2^, Fig. 2C), corresponding to radii ranging from about 25-35 μm (shapes appeared roughly spherical). These findings indicate that in the absence of Cx43 or with Cx43 lacking its carboxyl terminal domain, the blood brain barrier is more vulnerable to damage, although their infarct sizes are similar.

**Figure 2.**
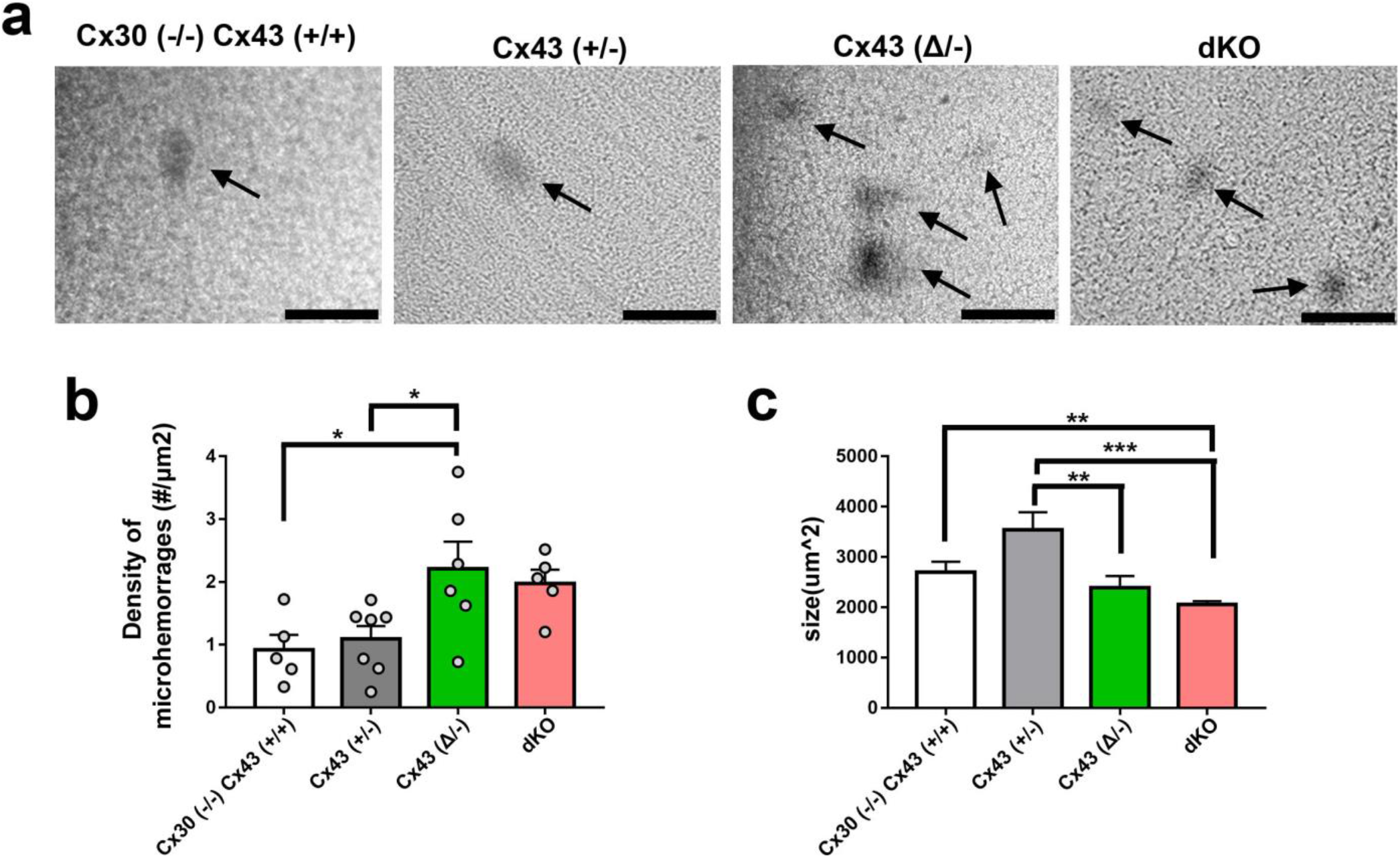
High pressure-induced brain leakage in Cx43 transgenic mice. (A) Bright-field images showing micro-hemorrhages (HRP leakage revealed with DAB in dark areas: arrows) in brains of mice as presented in Fig. 1 subjected to high hydrostatic pressure perfusion with a solution containing HRP. (B) Mean ± SEM values of the density of micro-hemorrhages measured in brains of mice of each genotype. (C) Quantification of the sizes of individual micro-hemorrhages in each genotype. Data from sections of whole hemispheres of 5 Cx43^(+/+)^/Cx30^(-/-)^, 6 Cx43^(+/-)^, 6 Cx43^(Δ/-)^ and 5 dKO mice. Data are presented as mean ± SEM (ANOVA followed by Tukey’s post hoc test : *p < 0.05, **p < 0.01, and ***p < 0.001.) Scale bars 100 µm.

### 3. Structured illumination microscopy (SIM) imaging of Cx43, AQP4 and AQP4ex in the perivascular domain

In order to compare resolution of conventional confocal images with those obtained using SIM, we focused on regions of the vessel wall at increasing magnification (Suppl. Fig. S1). In initial experiments, we immunostained for Cx43 in transgenic mice in which astrocytes were visualizable due to expression of yellow fluorescent protein (YFP) driven by GFAP promoter (mGFAP-Cre/RCE:loxP; RCE:loxP reporter mice (Sousa et al., 2009) bred with the mGFAP-Cre line (Garcia et al., 2004)) enabling us to distinguish astrocyte endfeet from cell bodies and from other components of the vascular wall (Suppl. Fig. S1A); resolution of Cx43 in the vessel wall was lost when the widefield image was zoomed to very high magnification (Suppl. Fig. S1B). When the widefield image in A was deconvolved and displayed at higher resolution, localization of Cx43 to astrocyte endfeet became apparent (Suppl. Fig. S1C). When Image in B was deconvolved using neighboring z-plane acquisition, a single gap junction plaque became resolvable (Suppl. Fig. S1D). SIM reconstruction of the image in D revealed further details of irregular topology, with subregions that were not visible without SIM in D (Suppl. Fig. S1E). Similarly, AQP4 puncta around vessels could be resolved in single plane widefield deconvolved images (Suppl. Figs S1F, G), where sub-200 nm details were observable using SIM (Suppl. Fig. S1H). The extended AQPex isoforms were also resolvable as small aggregates using isoform specific antibodies described in Methods (Suppl. Figs S1I - K). At highest magnification, 3D SIM revealed stripe-like patterns of particles (Suppl. Fig. S1K). As discussed below, AQP4ex isoforms have recently been shown to be located primarily in astrocyte endfeet, anchoring other isoforms ^21^.

### 4. Distribution of Cx43, AQP4 and AQP4ex in endfeet of transgenic mice with Cx43 mutations in astrocytes

SIM images of immunolabeling with Cx43 and AQP4 antibodies revealed the presence of both proteins in endfeet of each genotype, except that Cx43 was not detected in astrocytes in GFAPCre:”Cx43^fl/fl^ mice (Fig. 3). Figure 3A shows example images that correspond with data displayed in Figures 4 and 5, while Figure 3B shows example images for data in Figure 6. Although at this magnification differences among genotypes at the level of perivascular domains are not obvious (Figs 3A, B), 3D SIM analyses of Cx43 and AQP4 particle sizes revealed differences as described below.

**Figure 3.**
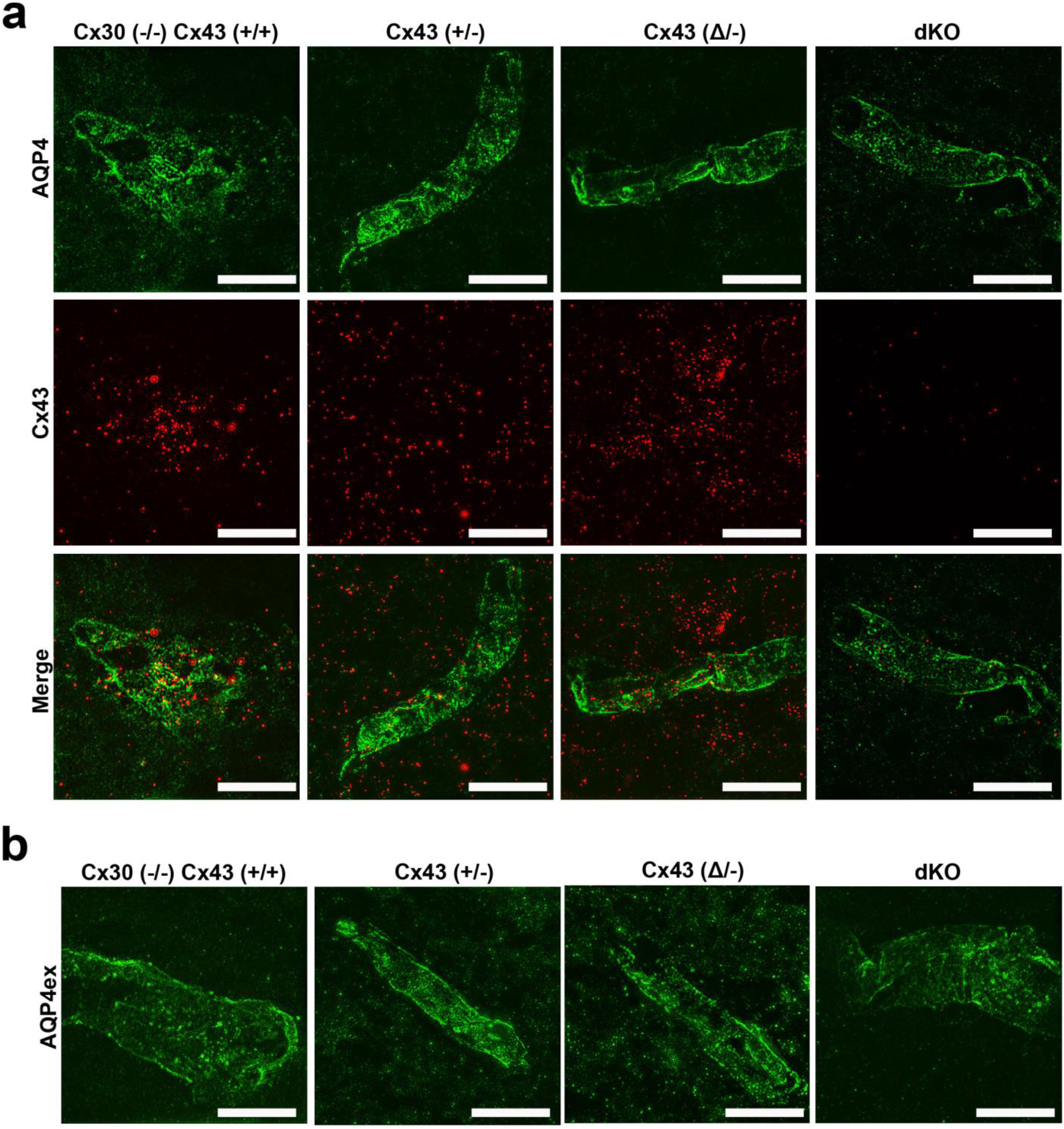
Distribution of Cx43, AQP4 and AQP4ex in brain sections from Cx43 transgenic mice. (A) 3D SIM reconstruction showing the expression of Cx43 (red) and AQP4 (green) in Cx43 ^(+/+)^/Cx30^(-/-)^, Cx43 ^+/-^, Cx43 ^(Δ/-)^, and dKO brain sections. (B) 3D SIM images showing the expression of AQP4ex (green) in brain sections from each genotype. Scale bars 10 µm.

**Figure 4.**
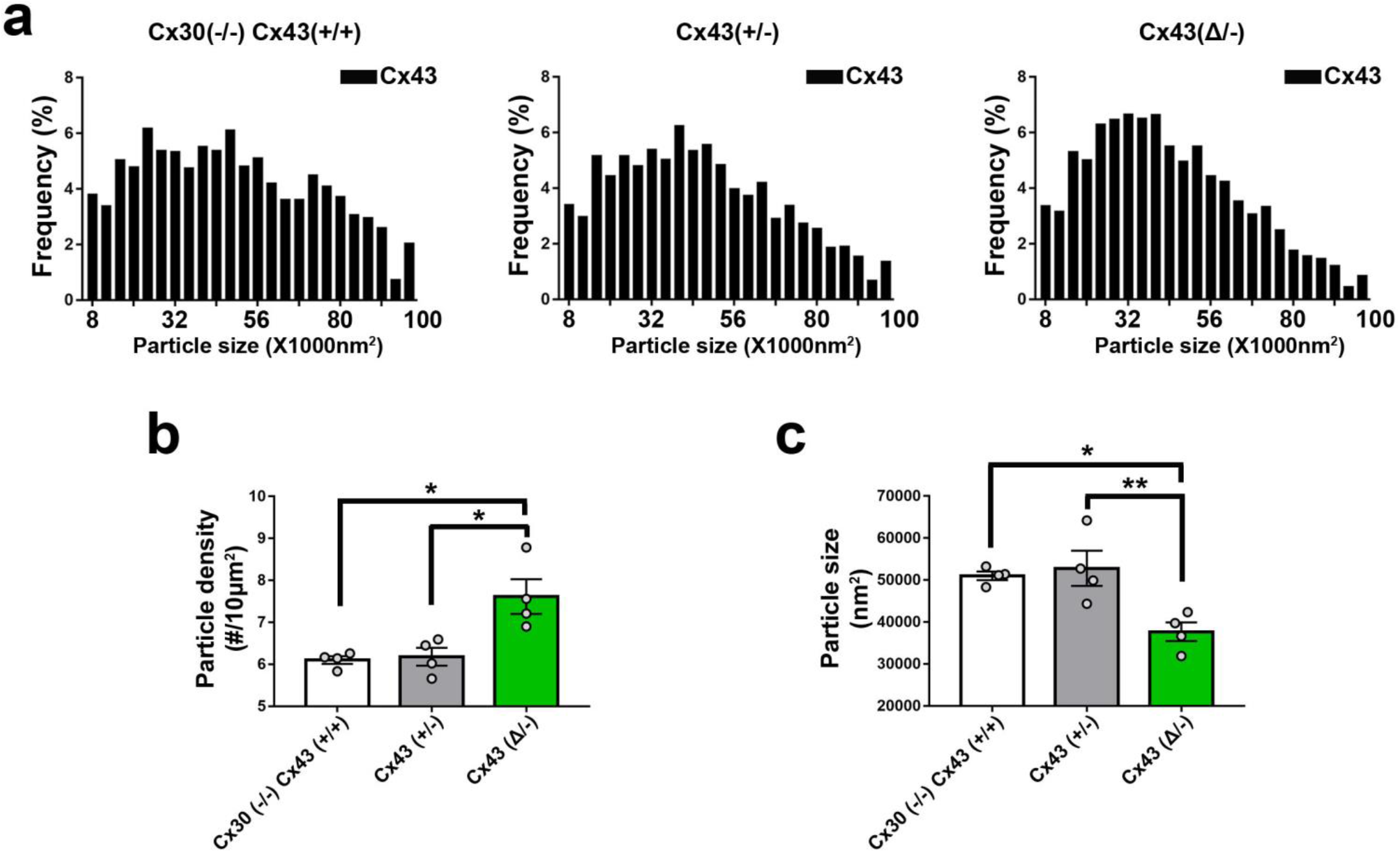
Distribution of Cx43 in endfeet of Cx43 transgenic mice. (A) Histograms showing size distribution of total number of particles obtained from 6-10 images of brain sections from each genotype (n = 4 mice, 1500-2000 particles analyzed from each genotype). (B) Comparison of Cx43 plaque densities for each of the transgenic lines (n = 4 mice, 6-10 images from each genotype). (C) Geometric means of particle sizes for perivascular Cx43 in each genotype (n = 4 mice, 6 - 10 images from each genotype). Data are presented as mean ± SEM (ANOVA followed by Tukey’s post hoc test: *p<0.05, **p<0.001.

**Figure 5.**
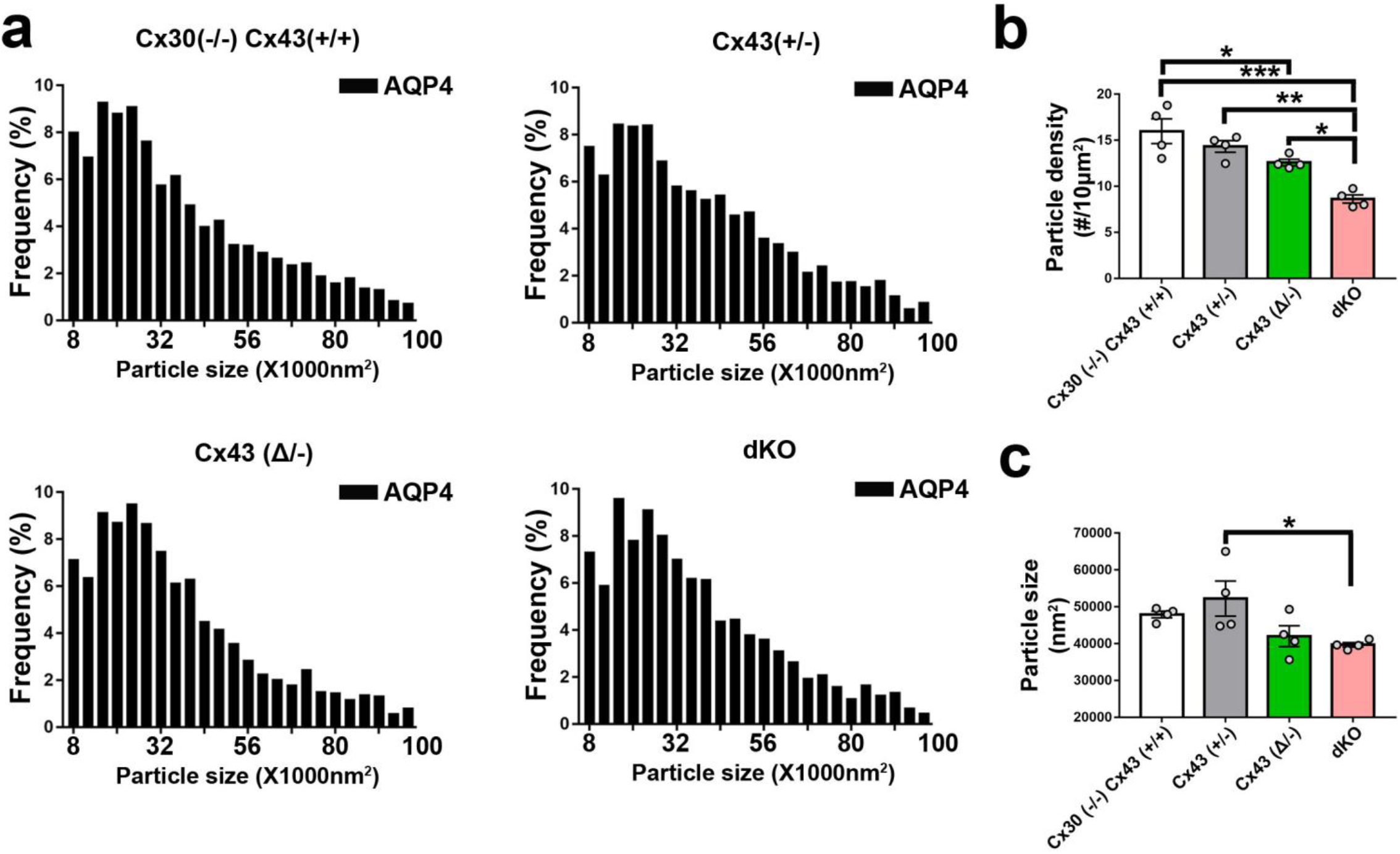
Distribution of AQP4 in endfeet of Cx43 transgenic mice. (A) Histograms showing size distributions of all AQP4 particles measured in each genotype. (n=4 mice, 6-10 images and -2000-3000 particles analyzed from each genotype). (B) Particle densities for AQP4 in perivascular domains from each genotype. Dots represent four individual brains, from which 6-10 sections were analyzed. Note that density of AQP4 particles in perivascular regions was negatively correlated with Cx43 expression in the various genotypes [Cx43^(+/+)^>Cx43^(+/-)^>Cx43^(Δ/-^)>Cx43^(-/-)^]. (C) Sizes of the particles were relatively constant among the genotypes, with only the Cx43^(+/-)^ differing from the dKO (n = 4 mice, 6 - 10 images analyzed from each genotype). Data are presented as mean ± SEM (ANOVA followed by Tukey’s post hoc test: *p < 0.05, **p < 0.01, and ***p < 0.001).

**Figure 6.**
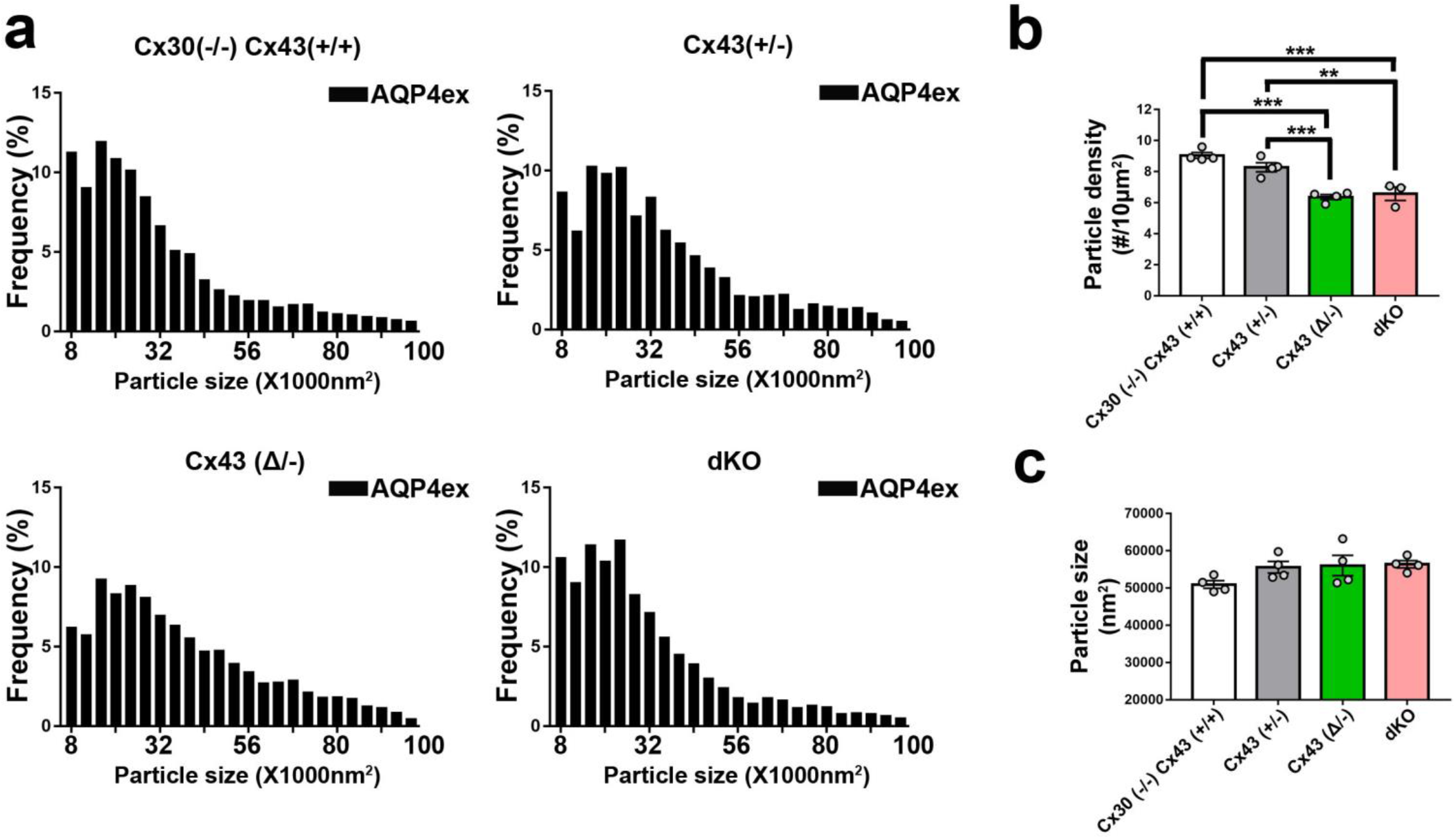
Expression of AQP4ex in astrocyte endfeet of transgenic mice. (A) Histograms showing size distributions of all AQP4ex particles measured in each genotype. (n = 4mice, 6 - 10 images and 2000 - 3000 particles analyzed from each genotype). (B) Particle densities for AQP4ex in perivascular domains from each genotype. Dots represent four individual brains, from which 6 - 10 sections were analyzed. Note that particle number was higher in WT and Cx43^(+/-)^ than in the truncation mutant and double KO. (C) Sizes of the particles were relatively constant among the genotypes. Data are presented as mean ± SEM (ANOVA followed by Tukey’s post hoc test: *p < 0.05, **p < 0.01, and ***p < 0.001.

#### a) Cx43 in endfeet of the genotypes

3D SIM as in Fig.3 allowed us to analyze sizes and numbers of Cx43 GJ plaques in the perivascular regions of brains of transgenic mice, with resolution of spots with areas less than about 8000 nm^2^ (corresponding to circles with radii ∼50 nm). Histograms of particle sizes showed distributions that were broadest for the Cx43 WT, intermediate with the heterozygote and narrowest in the truncation mutant (Fig. 4A). Mean Cx43 particle size was smallest for Cx43^(Δ/-)^ compared to Cx43^(+/+)^ and Cx43^(+/-)^ (Fig 4C). By contrast, Cx43 plaques were less abundant in Cx43^(+/+)^ and Cx43^(+/-)^ endfeet than for Cx43^(Δ/-)^ (Fig 4B).

#### b) Total AQP4 in endfeet of the genotypes

3D SIM of AQP4 particles allowed measurement of sizes in the same range as those for Cx43 (threshold radius <50 nm). Histograms (Fig. 5A) showed distributions more tightly clustered at smaller particle sizes than for Cx43 (Fig. 4A), although mean sizes were similar (∼50000 nm^2^; Fig. 5C). AQP4 particle density in Cx43^(+/+)^ was about twice that of Cx43 particles (1-1.5 *vs* 0.5-0.8/µm^2^) and declined in concert with Cx43 expression (Fig 5B): Cx43^(+/+)^>Cx43^(+/-)^>Cx43^(Δ/-)^>Cx43^(-/-)^. Sizes of the AQP4 particles was fairly constant across genotypes, although, as in case of Cx43, particles in the Cx43^(+/-)^ endfeet were largest (Fig. 5C).

#### c) Extended isoforms of AQP4 (AQP4ex)

3D SIM analysis of AQP4ex in the four Cx43 genotypes showed particle distribution histograms that were similar in overall topology to those of AQP4 (Fig. 6A). As was the case for total AQP4, mean particle density for AQP4ex was proportional to Cx43 expression (Fig. 6B). Sizes of particles were similar in all genotypes (Fig 6C).

#### d) Ratio of AQP4ex to total AQP4

In order to determine whether AQP4ex was preferentially localized in the perivascular region, we calculated the ratios of AQP4ex/total AQP4 in perivascular regions of each mutant. Our findings indicated that the ratio of AQP4ex/total AQP4 densities was highest in the Cx43^(-/-)^ and lowest in the Cx43^(+/+)^ (Fig. 7A). In contrast, the relative sizes of particles was negatively correlated with Cx43 expression, with Cx43^(+/+)^ having the lowest AQP4ex/AQP4 particle size ratio and Cx43^(-/-)^ having the highest ratio (Fig. 7B).

**Figure 7.**
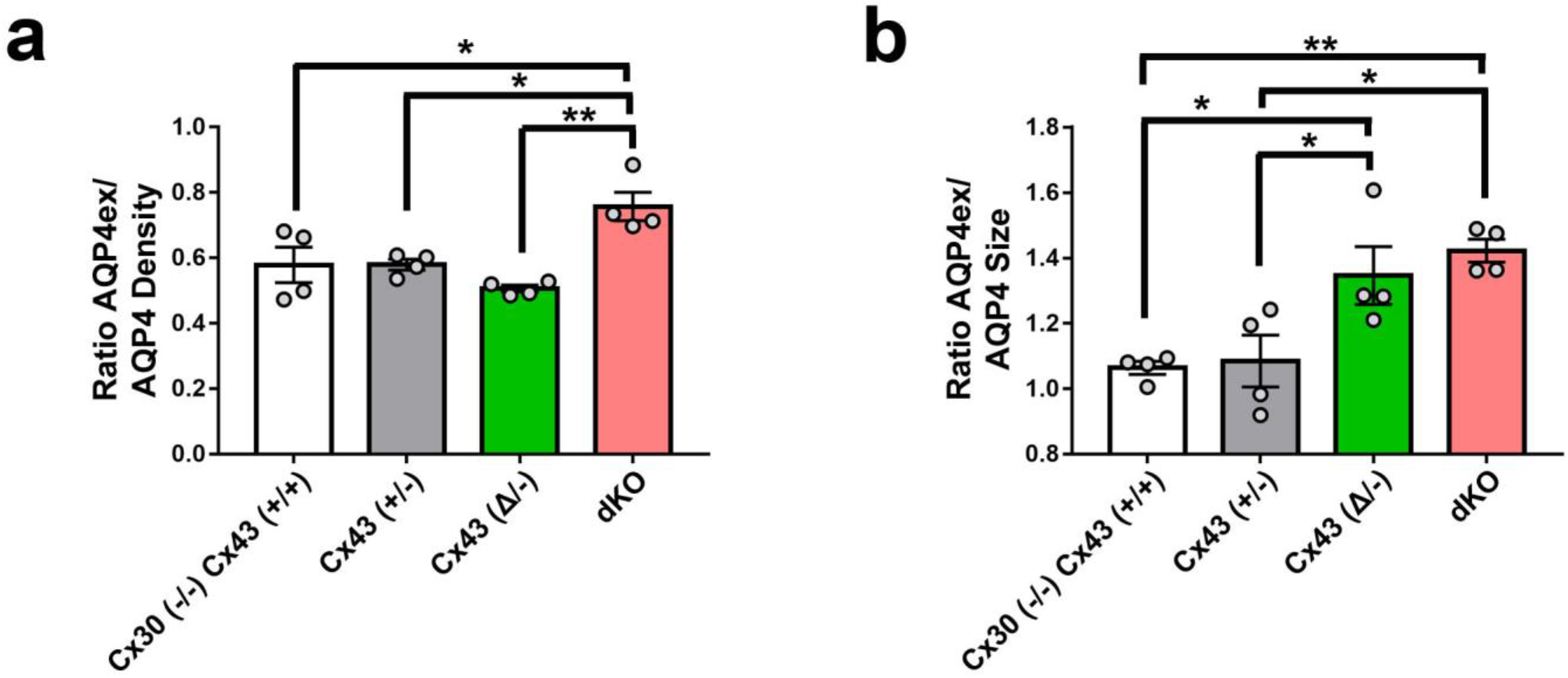
Ratio of AQP4ex/AQP4 density and size in perivascular brain regions of the Cx43 genotypes. (A) Ratios of particle densities of AQP4ex to AQP4 shows that dKO has higher ratio than the other genotypes. (B) Ratios of particle sizes shows that Cx43^(+/+)^ and Cx43^(+/-)^ are similar but the size of AQP4 is smaller than AQP4ex for the other groups. Data are presented as mean ± SEM (ANOVA followed by Tukey’s post hoc test: *p < 0.05 and **p <0.01).

### 5. Colocalization of Cx43 and Cx30 in perivascular astrocytes in Cx43^(Δ/-)^ and Cx43^(+/-)^ brain

We have shown in previous studies that Cx43 truncation leads to destabilization of GJ plaques in astrocytes ^6^, and we hypothesized that this would lead to remodeling of the endfoot domain with respect to other connexin proteins involved in function of the neurovascular unit. To test this hypothesis, we performed confocal microscopy on vascular domains in cortex. Fig. 8A shows representative fluorescence micrographs with dual labeling for Cx43 and Cx30. When overlap was quantified for the Cx43^(+/-)^ brain, we found that Mander’s coefficients for overlap of both Cx43 and Cx30 with total staining were about 0.2-0.35, while in the case of the truncation mutant (Cx43^(Δ/-)^) the coefficients were significantly higher (>0.6) (Fig. 8B).This finding indicates that the mobile Cx43 isoform is more highly interactive, consistent with our finding that the mobile mutants can readily penetrate GJ plaques ^7^.

**Figure 8:**
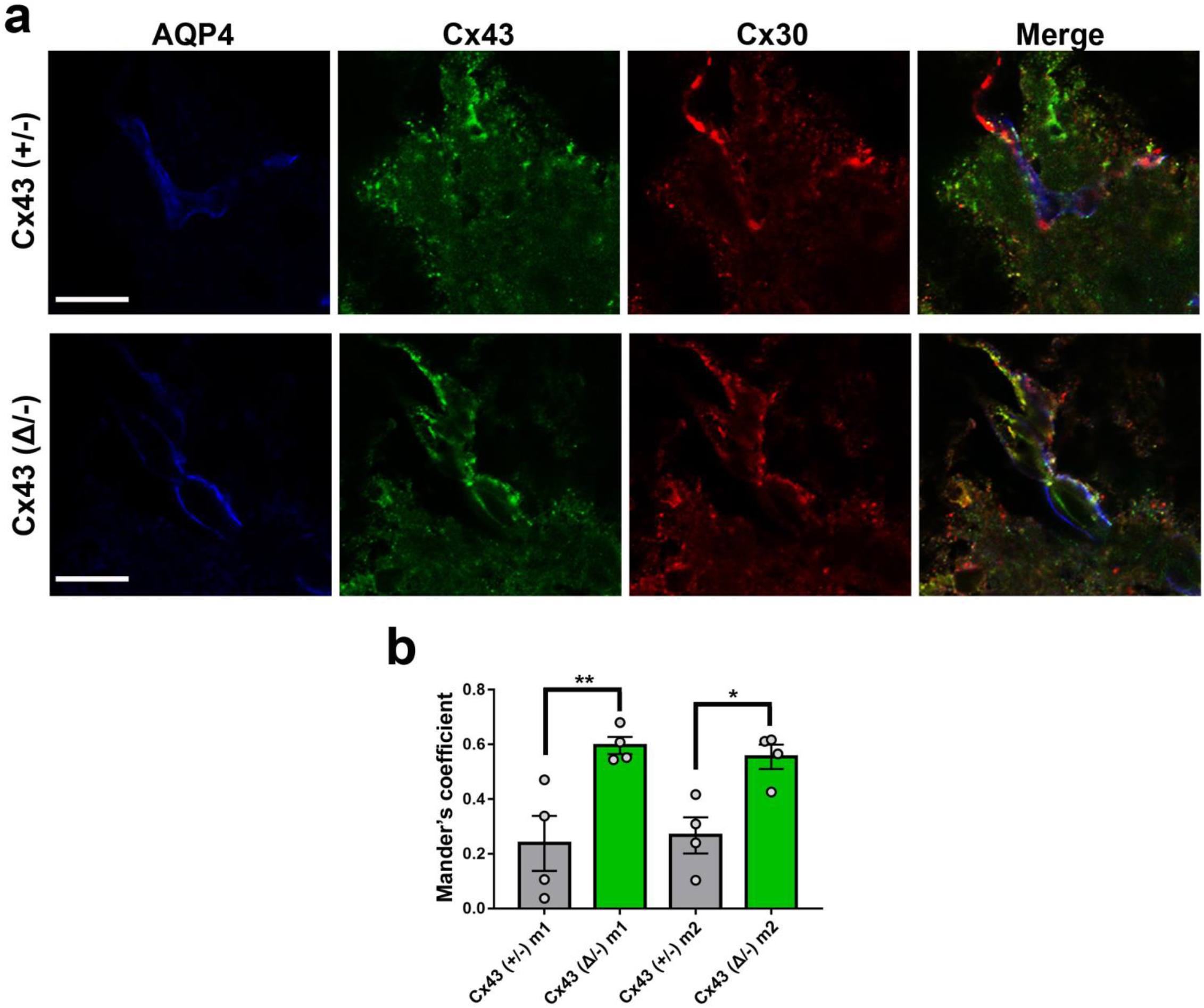
Higher co-localization of Cx43 and Cx30 in endfeet of Cx43^(Δ/-)^ than Cx43^(+/-)^ mouse brain. (A) Images showing expression of Cx43 (green) and Cx30 (red) in brain vessels of Cx43^(+/-)^ and Cx43^(Δ/-)^ mice. (B) Quantification of the degree of co-localization of Cx43 with Cx30 (m1) and of Cx30 with Cx43 (m2) at perivascular region of Cx43^(+/-)^ and Cx43^(□/-)^ mice. Note that overlap for each connexin with the other was higher in mice lacking the Cx43 carboxyl terminus. Scale bar 100 µm.

## DISCUSSION

In this study we have compared the perivascular organization of gap junctions and AQP4, the channels that are primarily responsible for water distribution in the astrocyte network, using transgenic mouse strains in which Cx43 abundance is reduced or its cytoplasmic binding domain was deleted. This perivascular structure, or glia limitans, is formed by astrocyte endfoot processes closely enwrapping the vessel wall. Both AQP4 and Cx43 are localized to astrocyte endfeet soon after birth (by P0 in mice), whereas Cx30 appears later (at about P12 in mouse) ^8,22,23^.

The four mouse genotypes examined here include those in which Cx43 expression is normal but Cx30 is deleted (Cx43^(+/+)^/Cx30^(-/-)^), Cx43^(+/-)^ or Cx43^(Δ/-)^ heterozygotes, where one allele is absent and the other is either WT Cx43 or one in which the coding region is exchanged with a truncated isoform (termed Cx43^(Δ/-)^ mice), and mice in which Cx30 is deleted and Cx43 ablation is targeted to astrocytes (Cx30^(-/-)^/GFAPCre::Cx43^fl/fl^, also termed dKO). Upon examination using endothelial CD31 immunostaining to localize distribution of Cx30, Cx43 and AQP4 within endfeet, we found that all vessels were macroscopically similar (Fig. 1), and immunolabeling was grossly similar except for absence of Cx43 in the GFAP::Cx43^fl/fl^ endfeet. Even higher magnification using 3D structured illumination microscopy (SIM) revealed no striking disorganization of the vessels or the endfeet (Fig. 3). This lack of gross alteration in vessel morphology is consistent with reports from others that brains were anatomically normal, with no gross difference in vascular morphology in the dKO (Cx30/GFAP::Cx43^fl/fl 8,9^ or in the Cx43^(Δ/-)^ mouse cortex ^13^.

Despite grossly normal vascular morphology, previous electron microscopy revealed that astrocyte endfeet were edematous in the dKO ^9 8^; moreover, oligodendrocytes were found to be vacuolated, which was interpreted as reflecting disruption of the gap junction mediated connections between astrocytes and oligodendrocytes ^9^, the so-called panglial syncytium ^24^. In our examination of HRP extravasation in the high-pressure perfusion model, we found that both Cx43 deletion and truncation increased vulnerability of the neurovascular unit to similar extent (Fig. 2), with number of micro-hemorrhages approximately double that in the respective controls. Sizes of most hemorrhages were similar, except that the Cx43^(+/-)^ brains showed somewhat larger areas of extravasation than the truncation brain and the infarct size in Cx43^(+/+)^ was larger than in the dKO. Therefore, these studies reveal that incidence of stroke vulnerability inversely correlates with Cx43 abundance and intact carboxyl terminal binding sites, conditions under which mobility and distribution of Cx43 and other nexus proteins are altered ^7^.

The initial astrocyte-targeted Cx43 knock-out mouse ^25^ displayed larger volume of injury after middle cerebral artery occlusion ^26^, accelerated spreading depression, and motor impairments ^25,27^. The residual coupling among astrocytes was abolished after crosses with a mouse deficient in the other major astrocyte connexin, Cx30 ^28^; while this has been taken as evidence that only Cx43 and Cx30 contribute substantially to astrocyte intercellular coupling, Cx26 is also normally expressed in at least in some astrocytes ^29^ and is syntonic with Cx30, so that Cx30 null mice also lack Cx26 expression ^30^. The Cx30-GFAPCre::Cx43^fl/fl^ mice (dKO) have depressed K^+^ buffering capacity and lowered threshold for epileptiform events ^28^, and they display more severe pathology when subjected to ischemia/reperfusion injury paradigms ^31^. Together, these studies provide extensive evidence that astrocyte connexins play an important role in maintaining ion homeostasis and minimizing damage in the CNS.

The Cx43 truncation mouse was generated in order to evaluate the role of the Cx43 carboxyl terminal domain in systems physiology. The surprising phenotype included pathologically elevated epidermal permeability, which has been attributed to loss of interaction with the tight junction protein ZO-1 ^10^. Studies on cardiac tissue from these mice have revealed higher propensity for ventricular arrhythmias during reperfusion and increased size and reduced number of GJ plaques in the myocardium (intercalated discs) ^12^. This result is the opposite of our finding of increased number and reduced GJ plaque size in astrocyte endfeet (Fig 4). This may call into question the widely accepted role of the carboxyl terminal PDZ binding domain in corralling Cx43, delimiting plaque size ^32^. Studies of the consequences of loss of the Cx43 carboxyl terminus in stroke reported that occlusion of the left coronary artery followed by 4 hr reperfusion led to more severe damage in Cx43^(Δ/-)^ than in the WT heterozygote hearts ^14^. In the brain, vessel permeation of India ink was similar in both Cx43^(+/+)^ and Cx43^(Δ/-)^ heterozygotes, but larger infarcts were seen in Cx43^(Δ/-)^ mouse brains when the middle cerebral artery occlusion model was performed ^13^.

Although previous studies have also shown that blood brain barrier vulnerability is higher in mice lacking astrocyte Cx43 or with the truncated construct, and that perivascular endfeet in these brains appear edematous, the organization of Cx43 in the astrocyte endfeet has not been examined. To determine more precisely how the expression of Cx43 and presence of its carboxyl terminal binding sites impact organization of proteins in the astrocyte endfoot we used 3D structured illumination microscopy (SIM) to achieve moderately high resolution of perivascular protein complexes in brain sections from these mice. Gap junctions can be most unambiguously detected in freeze fracture electron micrographs, where plaques containing even a small number of intramembrane particles can be identified, each of which is believed to be a gap junction channel, see ^33^. Histograms of particle sizes of Cx43 GJ plaques detected in our studies show minimal detectable size of about 8000 nm^2^, corresponding to a circular plaque with ∼50 nm radius that would contain ∼30 GJ channels within the plaque if interparticle spacing is ∼14 nm. When genotypes were compared, we found that density of perivascular Cx43 plaques was lowest in WT and Cx43^(+/-)^ mouse brains (∼6 compared to ∼8/10 µm^2^ in the truncation mouse), whereas sizes were smaller in the Cx43^(Δ/-)^ (∼0.23 vs ∼0.25 µm diameter) than in the other genotypes, so that there was little difference in total Cx43 at the endfoot (∼2.8 - 3.2% coverage of total perivascular area).

Like gap junctions, AQP4 forms supramolecular clusters decorating astrocyte plasma membranes. These so-called Orthogonal Arrays of Particles (OAPs) are formed by assemblies of AQP4 tetramers highly concentrated at the astrocyte endfeet. In freeze fracture studies, the lattice corresponds to 6.4 nm center to center spacing ^34-36^. Using antibodies recognizing all AQP4 isoforms or only AQP4ex, we achieved resolution of particle size comparable to that with Cx43, with smallest bin size of 8000 nm^2^ in the resulting histograms, similar to that reported by ^37^, corresponding to <500 square arrays. We found that in the perivascular endfeet, AQP4 particle density was directly proportional to Cx43 expression level (Cx43^(+/+)^>Cx43^(+/-)^>Cx43^(Δ/-)^>dKO). Particle sizes were slightly higher for Cx43^(+/-)^ heterozygotes and lowest for dKO. Coverage of the endfoot was considerably higher for Cx43^(+/+)^ (∼7.5%) than for the others and was lowest in the dKO (∼3.2%). Compared to Cx43 plaques, AQP4 particles were about 50% more abundant but similar in size.

AQP4 is distinctive among aquaporin water channels in its expression as several distinct isoforms through two mechanisms that modulate protein translation: Leaky scanning, in which a weak initiation codon may be skipped, generating alternate M1 and M23 AQP4 isoforms ^35^, and translational read-through, in which translation continues beyond a stop codon, resulting in longer N-termini, termed AQP4ex to denote the extension ^21,38^. Using antibodies recognizing only the AQP4ex isoforms, we determined distribution of particles in the perivascular region and were able to compare to overall AQP4. As in the Cx43 plaque and AQP4 OAP studies, we achieved a resolution limit of 8000 nm^2^. AQP4ex density was found to be higher in Cx43^(+/+)^ and Cx43^(+/-)^ than in Cx43^(Δ/-)^ and the dKO, whereas particle sizes were similar in each group, so that endfoot coverage was slightly lower in the dKO than in WT (ranging from about 3.5-4.5%). Sizes of particles formed of AQP4ex were previously reported to be larger than for total AQP4 ^39^, which was confirmed in our measurements in dKO and Cx43^(Δ/-)^ but not in the Cx43^(+/+)^ or Cx43^(+/-)^ groups.

Orthogonal particle arrays have long been known to be present in astrocyte membranes ^40^ and have been used for identification of astrocytes in freeze fracture replicas. AQP4 isoforms comprise these OAPs and show profound polarization in expression at the endfeet ^41-43^. The polarized distribution has been attributed to the presence of scaffolding molecules to which AQP4 binds with high affinity, primarily dystrophin/syntrophin (see ^44^), and one role of the readthrough extension of AQP4ex may be to anchor this isoform in the endfoot through interactions with syntrophin and other scaffolds ^21^. Recent studies using mice in which AQP4ex or syntrophin isoforms were deleted have revealed that AQPex expression appears to reciprocally localize α-syntophin at the endfoot and that deletion of both syntrophin isoforms leads to loss of AQP4 at the endfoot ^21,45^. An additional finding from the study on syntrophin deletion was that endfoot AQP4 OAPs were smaller, while Cx43 plaques increased in size ^45^. In addition to preferential adhesion to scaffolding proteins as a mechanism that enhances endfoot expression of AQP4 and especially the AQP4 isoform, it has recently been reported that local translation of certain astrocyte proteins occurs in endfeet ^46^. Regulatory proteins that direct translation of individual AQP4 isoforms have been discovered (see ^47^), raising the possibility that local translation may bias expression in specific cellular compartments. Combined with selective retention by cytoskeleton-linked scaffolds, this could lead to both stable and dynamic polarization of the endfoot to respond to local demands for water flux.

The perivascular endfoot gap junction plaques include both Cx30 and Cx43, as evidenced by cryo-immunogold electron microscopy ^48^. Although FRAP studies indicate that these connexins can intermix in transfected cells ^7^, there are no reports indicating whether these channels are segregated or intermixed *in vivo*. To test this hypothesis, we used super-resolution microscopy to determine in brain tissue whether Cx30 and Cx43 are co-localized within plaques in astrocyte endfeet and whether co-localization differed in the vascular unit in the cerebellum, where Cx30 is reportedly higher than in cortex or hippocampus ^49^ or in thalamus, where Cx43 expression is rather low ^50^. Analysis of colocalization of Cx30 with truncated Cx43 (Cx43^(Δ/-)^) in perivascular domains in these mice revealed much higher colocalization than in the Cx43^(+/-)^ controls. This result is consistent with our photobleach recovery experiments, which showed that Cx30 is much more penetrant into Cx43^(Δ/-)^ GJ plaques than into Cx43^(+/+)^ GJs ^7^.

Together, AQP4 and GJ channels connect astrocytes into a water-equilibrating network, where the GJ-connected glia extends from arteriolar astrocyte endfeet that take up water, glucose and other solutes, across brain parenchyma, where water balance is maintained and energy substrates are delivered, to venous perivascular astrocyte endfeet where efflux of excess water and metabolites occurs into the circulation. Location, expression and function of AQP4 and GJs are interdependent, as indicated by our initial studies of AQP4 siRNA effects on Cx43 in astrocytes ^51^ and confirmed by more recent examination of transgenic mice in which either AQP4 ^4,45^,52 or Cx43 ^53^ is manipulated. Such interdependence likely provides a major mechanism by which water and solute homeostasis is achieved, and disruption of normal distribution of these two proteins impacts brain edema and BBB permeability ^54^.

Interestingly, OAP alterations have been observed in different pathological conditions such as brain tumors, ischemia and Alzheimer’s disease ^55-57^. Moreover, the interest in OAP studies has recently increased since the discovery of their involvement in the pathogenesis of Neuromyelitis Optica (NMO), a CNS autoimmune channelopathy whose autoantibodies are only able to attack their antigen AQP4 if arranged in OAPs ^58-60^. Astrocyte endfeet edema has been observed in association with BBB leakage in the dystrophic MDX mouse model ^61^ during hypoxia ^62^ or stroke ^63^. Thus, in dKO mice, it may directly affect the endothelial barrier properties as well. In particular, the regulations taking place in astroglial endfeet inducing BBB phenotype might be severely compromised.

In conclusion, these functional and super-resolution imaging studies of perivascular regions of transgenic mice support the hypothesis that Cx43 and its carboxyl terminal domain contribute to structural organization of the astrocyte endfoot. In the absence of Cx43, or when its carboxyl terminus is lacking, the blood brain barrier is more fragile and sizes and distribution of AQP4 and its extended isoform are altered. Thus, as in cultured cells where exogenous expression of fluorescent Cx43 isoforms has revealed impact of its carboxyl terminus on lateral mobility and interaction with other proteins^7^, immobile Cx43 in gap junction plaques between endfeet appears to provide a framework that contributes to establishing and maintaining polarity of astrocyte processes.

## MATERIALS and METHODS

### Animals

Cx43 genotypes of mice used in these studies are illustrated in Fig 1. They include Cx43^(+/+)^/Cx30^(-/-)^, Cx43^(+/-)^, Cx43^(K258stop/-)^, and Cx30^(-/-)^/hGFAP-Cre:*Cx43*^*fl/fl*^ (termed respectively Cx43^(+/+)^, Cx43^(+/-)^, Cx43^(Δ/-)^ and dKO). The dKO and the Cx43^(+/+)^/Cx30^(-/-)^ littermates (about 6 months old males and females) were maintained at the Experimental Therapy Unit, University Clinical Center Bonn, Germany. Maintenance and handling of animals was according to local and government regulations. Mice were kept under standard housing conditions (12 hr/12 hr dark–light cycle, food, and water ad libitum). Experiments were approved by the North Rhine– Westphalia State Agency for Nature, Environment and Consumer Protection. Mice expressing Cx43 with a stop codon at aa 258 (Cx43^K258/^; referred here as Cx43^(Δ/-)^) were generated by replacing Cx43 with Cx43^K258stop^ ^10^. Because homozygous Cx43^(Δ/Δ)^ mice are not viable, Cx43 heterozygous null (Cx43^+/-^) and Cx43^Δ/-^ mice maintained at NYU College of Medicine animal facilities and NYIT College of Osteopathic Medicine animal facilities were bred to obtain Cx43^(Δ/−)^ and their controls, Cx43^(+/-)^ mice. Six to 12 months old male and female mice of these genotypes in a mixed background (129/Ola/C57BL/6) were chosen after litter genotyping performed as previously described ^10,12^. All procedures were approved by respective institutional IACUC.

### BBB permeability assay

For *in vivo* brain high hydrostatic pressure perfusion (HPP), we used a solution which included albumin as an agent to increase viscosity/hydrostatic pressure [^8^; method derived from ^17^ and modified to include elevated hydrostatic pressure ^18^]. In brief, mice were anesthetized with a cocktail of xylazine (20 mg/kg)-ketamine (100 mg/kg); deeply anesthetized animals were intraperitoneally (i.p.) injected with norepinephrine (NE: 20 µg/kg) and 5 min later, the thoracic cavity opened to expose the heart for perfusion. The right ventricle was severed and a horseradish peroxidase (HRP: 3.5 mg/ml) Krebs bicarbonate buffered solution containing NE (2 µg/kg) and serum albumin (40 mg/ml) was perfused for 2 min into the left ventricle using a syringe pump at a rate of 2.5 ml/min. This was followed by a 2 min perfusion with HRP-Krebs bicarbonate buffered solution (albumin- and NE-free) and then by a 4 min perfusion with 4% paraformaldehyde solution. Brains were removed and post-fixed for 1 hr in 4% PFA prior to cryopreservation in serial sucrose (10 - 30%) solutions. OCT-embedded brains from four Cx43^(+/+)^/Cx30^(-/-)^, four dKO, six Cx43^(Δ/−)^ and six Cx43^(+/−)^ mice were cryosectioned for HRP staining (using DAB) and for Cx43, Cx30, AQP4 and AQPex immunohistochemistry.

### Peroxidase staining and image analysis

To localize peroxidase activity in HRP perfused brains, the Sigma Fast DAB (3,3′-diaminobenzidine tetrahydrochloride; Sigma-Aldrich) reagents were used according to the manufacturer’s instructions. Briefly, one DAB tablet and one urea hydrogen peroxide tablet were dissolved in 14.5 ml ultrapure water and 0.5 ml enhancer (0.3% CoCl_2_) solution added to the DAB reaction solution. Slides containing brain sections were incubated in DAB reaction solution for several min until color developed. Reaction was stopped by washing the slides in PBS which were then mounted in mounting medium for analysis. Bright-field images of 5 Cx43^(+/+)^/Cx30^(-/-)^ KO, 5 dKO, 6 Cx43^(+/-)^ and 5 Cx43^(Δ/-)^ brains stained with DAB were acquired using an inverted Nikon microscope equipped with 4x objective and Metafluor software. Brain section images were evaluated for possible differences in the number, size and density of micro-hemorrhages (DAB positive areas) among the genotypes. For each mouse, quantification was performed using ImageJ software ^19,20^ to measure area infiltrated and density of microhemorrhages in DAB-reacted sections on a complete set of sections throughout a brain hemisphere.

### Immunohistochemistry

Cryosections of HHP mouse brains were stained for Cx43, Cx30, AQP4, and its extended isoform AQP4ex. Cellular distributions were examined using confocal imaging; particle sizes and densities near brain vessels were determined using super-resolution microscopy (structured illumination microscopy: SIM). Briefly, isolated tissue was post-fixed in 4% PFA solution overnight at 4°C, then washed in PBS, immersed in 30% sucrose solution overnight and finally embedded in Tissue OCT compound (Leica, Wetzlar, Germany). Sections of 10 μm thickness were cut on a cryostat (CM 1900; Leica, Wetzlar, Germany) at -20 °C.

After incubation with blocking solution (PBS containing 2% BSA and 0.4% Triton-X100), brain cryosections were incubated overnight at 4°C with either mouse anti-Cx43 (1:300; Millipore, cat# 05763) and rabbit anti-AQP4 (1:500; Sigma cat# A5971) antibodies or with mouse anti-Cx43 (1:300; Millipore, cat# 05763) and rabbit anti-Cx30 (1:500; Life Technologies, cat# 71-2200) diluted in blocking solution. In addition, we used a custom rabbit polyclonal anti-mouse AQP4ex (a gift from Drs Frigeri and Nicchia) generated against the peptide DSTEGRRDSLDLASC within the mouse AQP4 carboxyl terminal extension (GenScript Biotech, Piscataway, NJ, USA) that has been shown to detect the extended AQP4 isoforms ^21^. To stain vessel walls, we used antibody goat anti-CD31 [platelet endothelial cell adhesion molecule, PECAM-1(1:100; Santa Cruz Biotechnology; cat# M-20, sc-1506)]. After several washes in PBS-T, tissues were incubated for 1 hr at RT with Alexa-Fluor 488 goat anti-mouse (Invitrogen cat# A11012), Alexa-Fluor 488 donkey anti-rabbit (Invitrogen cat# A-21206), Alexa-Fluor 594 goat anti-rabbit secondary antibodies (Life Technologies, cat# 11001) and Alexa-Fluor 488 donkey anti-goat (Invitrogen cat#A11055) diluted 1:1000 in phosphate buffered saline with 0.1% Tween 20 detergent (PBS-T). After washes in PBS, slides with brain sections were mounted for microscopy using ProLong Diamond anti-fade mounting medium with DAPI (Molecular Probes, Cat# P36971) to stain cell nuclei.

### Imaging acquisition

For confocal microscopy, a Leica inverted DMi8 SP8 microscope and Leica LASX software were used for image acquisition and analysis (Leica Microsystems CMS GmbH). A 63X NA=1.4 Oil PLAPO, WD=0.14mm WHITE objective immersion liquid with a refractive index of 1.5 was used. All confocal images were collected using 594 and 488 nm lasers for excitation and a pinhole diameter of 1 Airy unit. A minimum of four ROIs were acquired per slice and processed for analysis using Image-J.

### Structured Illumination imaging (SIM)

For SIM, images of Cx30, Cx43 and AQP4 and its extended AQP4ex isoforms in immunostained brain sections were acquired from each genotype and analyzed using a Nikon ECLIPSE Ti-E microscope and Nikon Elements software, equipped with a oil-immersion Nikon 100X widefield Objective with a refractive index of 1.5, and a Zeiss Elyra S1 with 63X NA1.4 Objective, with 5 grating positions and 40 nm pixels on an EMCCD camera were used. Excitation of the Alexa Fluor488 and Alexa Fluor594 dyes was achieved with 488 and 561 nm laser beams, and the emitted fluorescence was collected through GFP and RFP emission filters, respectively, using an EMCCD camera (Andor iXon3 DU897).

### Image analysis

To assess the expression of Cx43, AQP4 and AQP4ex particles localized along blood vessels in the brain cortex, SIM images were acquired from perivascular regions selected randomly within each section. The number and sizes of the particles were determined from mice of each genotype using a plugin (“Analyze Particles”) of ImageJ. For that, SIM stacked images were converted to monochrome (8-bit) images, thresholded and particles outlined. The number and size (µm^2^) of outlined Cx43 and AQP4 particles obtained for each set of images were then exported and statistically analyzed in GraphPad Prism6 software.

As is illustrated In Suppl. Fig. 2, log-transformed values of particle sizes were normally distributed; means of log-transformed data or geometric means of non-transformed data sets were used to compare particle sizes and relative density (particles/µm^2^).

### Colocalization of membrane proteins

To evaluate the degree of co-localization of Cx43 and Cx30 puncta at the perivascular domains, brain sections of Cx43^(+/-)^ and Cx43^(Δ/-)^ mice were immunostained with anti Cx43 and anti-Cx30 antibodies. Z-stack images were acquired using a Leica inverted DMi8 SP8 microscope confocal microscope and images analyzed using a plugin (“Co-localization Threshold”) of Image-J (NIH-Fiji). For that, regions of interest (ROIs) were placed on each Cx43 immunofluorescent spot located along a blood vessel after splitting the two merged channels (488 and 594 nm). The Mander’s correlation coefficient with Costes’ auto-threshold detection was used to estimate the fraction of Cx30 fluorescence (594 nm channel) that co-localized with Cx43 (488 nm channel) (M1) and the fraction of Cx43 that colocalized with Cx30 (M2) within the ROIs throughout the z-stacked images.

### Statistical analyses

Results are presented as the mean ± SEM. Shapiro–Wilk normality test was performed to evaluate the data distribution. Statistical significance was evaluated using unpaired Student’s t test or ANOVA followed by Tukey’s post hoc with multiple statistical comparisons between groups using GraphPad Prism 7 software. Differences were defined as follows: *p < 0.05, **p < 0.01, and ***p < 0.001.

## Acknowledgements

Grant support: R21NS116892 and NS092466 (DCS and ES), EU project H2020-MSCA-ITN 722053 EU-GliaPhD (CS), BMBF projects 16GW0182 CONNEXIN and 01DN20001 CONNEX (CS). We gratefully acknowledge Drs Antonio Frigeri and Grazia Paola Nicchia (University of Bari, Italy) for the gift of anti-AQP4ext antibody, the technical assistance of Jian Pan with cryosections, and Dr. Raddy Ramos assistance with Cx43^(Δ/-)^ colony establishment and breeding at the animal facility at New York Institute of Technology College of Osteopathic Medicine.

## Authors Contribution

DCS and ES conceptualize the study; KM, GS, CS, PG helped setting up the experiments; AC, RS, AT, LM, PG, ES performed experiments; AC, RS, ES, DCS analyzed the data; DCS lead the writing in consultation with AC, RS, ES; KM, GS, KS provided critical feedback and helped writing the paper.

## Additional information

Supplementary information: Supplementary Figures S1 and S2.

**Correspondence and requests for materials** should be addressed to David C. Spray or Eliana Scemes.

