## Supplemental Figures S1 and S2 for "Cx43 carboxyl terminal domain determines AQP4 and Cx30 endfoot organization and blood brain barrier permeability"

### SUPPLEMENTARY MATERIAL

**Supplementary Figure 1. Super-resolution microscopy of Cx43 (A-E), AQP4 (F-H), and AQP4ex in the astrocyte endfoot in brain sections.** (A) Confocal 3D reconstruction from GFAP-YFP mouse showing cytoplasmic YFP in astrocyte cell body and endfeet (green) and Cx43 immunostaining (red). (B) Zoomed in image of single plane, Cx43 (red) channel only showing lack of sub-500 nm resolution with this confocal microscope and objective/detector. (C) Deconvolved widefield image of a vessel wall at higher magnification than in A. (D) Zoomed image of Cx43 staining from B in widefield with deconvolution (using neighboring z-plane acquisition). (E) SIM reconstruction of the same plaque, note plaque shape and sub-regions can be resolved in E that cannot be resolved in D. (F) Deconvolved single plane widefield image of AQP4 in astrocyte endfeet wrapped around a vessel running diagonally across the image. (G) Inset of the deconvolved immunostained image. (H) Structured Illumination Microscopy reconstruction of the inset region reveals resolution of sub-200 nm features; white arrows indicate portions of the AQP4 signal that can only be recognized with super-resolution imaging parameters. Imaging performed on a Zeiss Elyra S1 with 63X NA1.4 Objective, with 5 grating positions and 40 nm pixels on an EMCCD camera. (G) Super-resolution microscopy of AQP4ex in the astrocyte endfoot in brain sections, illustrated at increasingly higher magnification. (I) Confocal 3D reconstruction showing the expression of AQP4ex (green) at the perivascular level in Cx43$^{(+/-)}$ mouse brain sections. Scale bar: 50µm. (J) 3D SIM reconstruction of AQP4ex (green) showing the same vessel of I (white box). Super-resolution imaging allows insights into structural features of the AQP4ex at perivascular level. Scale bar 10 µm. (K) Zoomed 3D SIM image (from J, white box) showing at higher magnification the AQP4ex particles at perivascular level. Scale bar 3 µm.

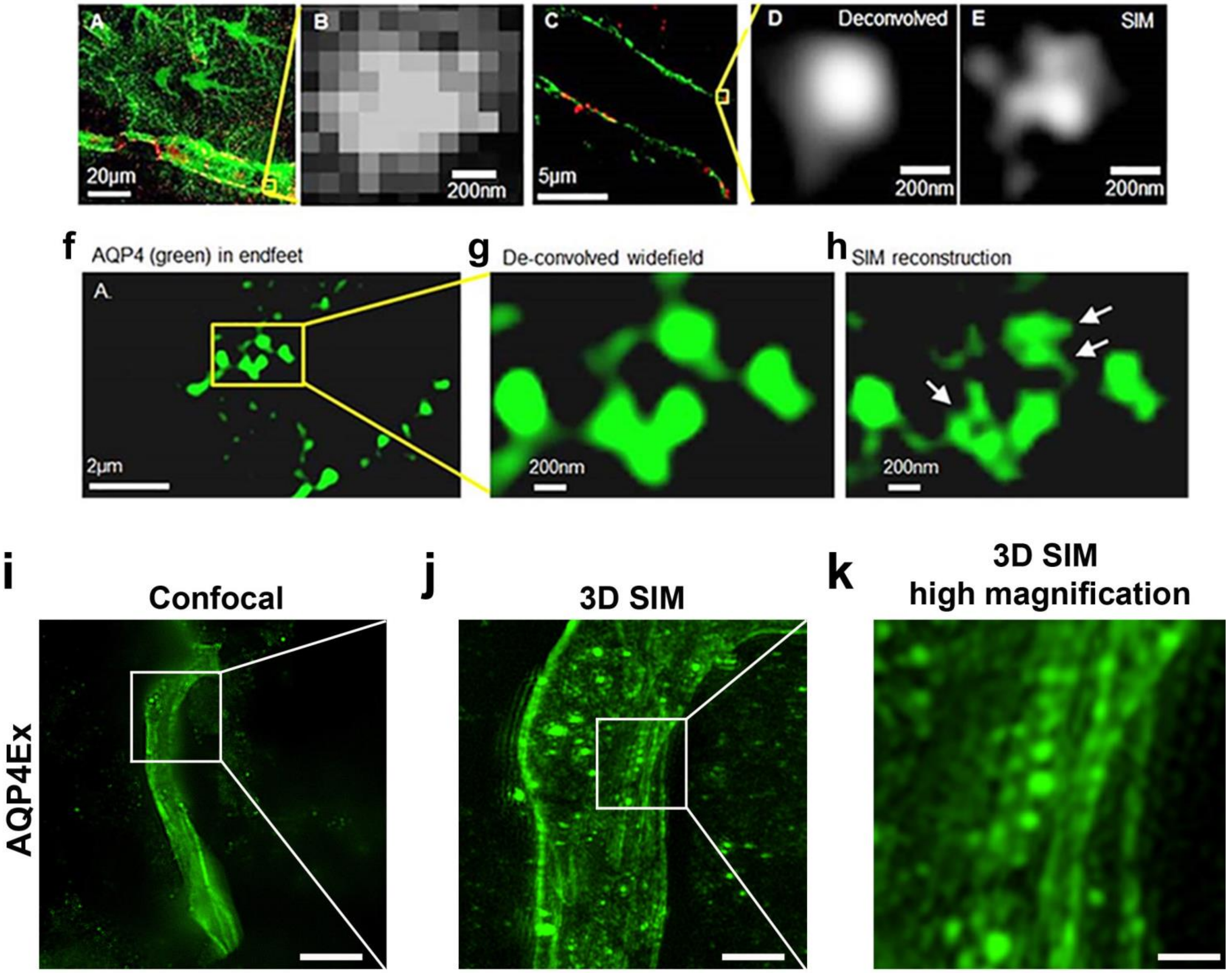
A
20μm
B
200nm
C
5μm
D
Deconvolved
200nm
E
SIM
200nm
f
AQP4 (green) in endfeet
A.
2μm
g
De-convolved widefield
200nm
h
SIM reconstruction
200nm
i
Confocal
j
3D SIM
k
3D SIM
high magnification
AQP4Ex

**Supplementary Figure 2. Log transformed histograms to illustrate method of counting particles.** In order to determine whether particle sizes were normally distributed, we log normalized all data a plotted in histograms. (Shapiro–Wilk normality test ***p < 0.001).

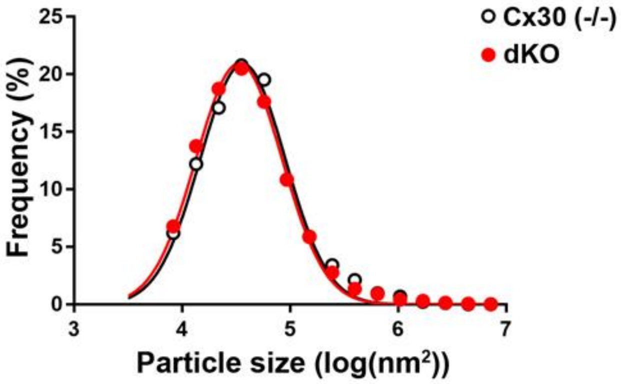
